# Early visual experience contributes to the calibration of higher-order body representations

**DOI:** 10.64898/2026.11.11.766044

**Authors:** Yuqi Liu, Yifei Wang, Yueyue Ma

## Abstract

Higher-order body representations, including representations of body shape and size, play an important role in maintaining perceptual constancy in touch. However, whether the development of these representations depends on early visual experience remains unclear. Here, we investigated tactile distance perception in congenitally blind, late-blind, and sighted individuals. A given tactile distance is typically perceived as longer on the finger than on the forearm, reflecting distortions in cortical topography that is partially rescaled by higher-order representations of body size and shape. Congenitally blind participants showed a greater tendency to perceive identical tactile distances on the index finger as longer than those on the forearm. A similar bias was observed when comparing the forehead and face, suggesting that altered tactile distance perception in congenital blindness is not limited to body regions with extensive sensory-motor experience. Furthermore, within the late-blind group, the similarity of perceptual patterns to sighted controls, relative to congenitally blind participants, was associated with blindness duration. Together, these findings suggest that both early visual and post-blindness experience contributes to the calibration of tactile distance perception across the body, with its influence potentially extending beyond body regions extensively engaged in sensory-motor activities.

## Introduction

Accurately representing tactile information on the body is crucial for sensing and interacting with the environment. While the primary somatosensory cortex is largely organized somatotopically (S1), with neighboring skin areas represented by neighboring cortical regions, additional higher-order representations have been proposed to which information from S1 is transformed and eventually to tactile percepts (Medina & Coslett, 2010; Longo et al., 2010). One of the representations concerns body shape and size (“body form representation”, Medina & Coslett, 2010; Longo et al., 2010; de Vignemont, 2010; Schwoebel & Coslett, 2005). For example, in individuals with lesions involving S1, tactile localization preserved the spatial topography of the hand despite systematic mislocalization of touch, suggesting a representation that encodes the relative location on the hand, i.e. the hand form (Rapp et al., 2002).

The body form representation plays an important role in processing tactile distance and size (Longo et al., 2010; de Vignemont et al., 2005). When two tactile stimuli are applied, their corresponding neural correlates would be activated. However, neural activation itself does not directly inform the distance between touch in the physical world. To map S1 activation to distance, the brain needs to reference the size and shape of the body. In a form of Weber’s illusion, two tactile points of fixed distance were perceived as further apart when they were applied to the finger than to the forearm, reflecting a higher receptor density in the finger area than the forearm area in S1 (cortical magnification, Taylor-Clarke et al., 2004, de Vignemont et al., 2005). Crucially, the difference in perception is disproportionately smaller than the difference in receptor density (30% vs. 340%, Green 1982), suggesting that information from S1 is corrected by a representation of the body size and shape, a mechanism that may contribute to maintaining perceptual constancy across different skin regions.

One question concerns the nature of the body form representation. Past studies reported altered perceived hand size after visual exposure to a larger or smaller hand, indicating that visual information contributes to the body form representation. In Taylor-Clarke et al. (2004), simply viewing an expanded forearm and a shrunk hand reduced the difference in the perceived tactile distance between these skin areas, indicating the causal role of vision on perceived body size and in turn, tactile distance perception. Beyond tactile size perception, online visual information modulates various aspects of tactile processing including tactile acuity (Longo et al., 2008; Cardini et al., 2012), sensitivity (Ro et al., 2004), localization (Margolis & Longo, 2015; Liu & Medina, 2021), and pain perception (Moseley et al., 2008), suggesting a general role of vision in modulating body representation. On the other hand, perception of body size can be manipulated proprioceptively (e.g. the Pinocchio illusion, Lackner, 1988; de Vignemont et al., 2005; Ehrsson et al., 2005) or with anesthesia (Gandevia & Phegan, 1999; Paqueron et al., 2003), when individuals were blindfolded. In other cases, transient changes in the perceived body form occurred despite invariant visual inputs (Podoll & Robinson, 2000, 2002). These findings indicate that the body form representation is of multisensory nature, being informed by multiple sensory modalities and relying on the integration across modalities.

While in typical adults the body form representation is multisensory, its developmental origin remains unaddressed. Specifically, does it require early visual experience to form a near-veridical representation of one’s body size and shape? On one hand, visual information may play a particularly important role during the initial formation of body form representations (Thaler et al., 2018; Martel et al., 2016; Buxbaum & Coslett, 2001; Schwoebel & Coslett, 2005). By providing direct information about the size and shape of the body in relation to the external world, vision may help establish a metric representation of the body that is calibrated to the physical dimensions of objects in the environment. In the absence of early visual experience, such an external metric reference may be lacking, potentially resulting in a less consistently calibrated representation of body size. Consistent with this hypothesis, congenitally blind children built substantially distorted body figures compared with typically-developed children (Kinsbourne & Lempert, 1980). On the other hand, individuals with congenital blindness have been shown to exhibit superior performance in object size estimation (Smith et al., 2005), suggesting that alternative mechanisms based on haptic exploration and sensorimotor interactions can support the development of metric representations of object size in the absence of vision.

Recent work has begun to examine the representation of body metrics in congenital blindness, yielding inconsistent findings. One study reported magnified distortion in the hand model as measured with the landmark localization task (Kottu & Lazar, 2025; Longo & Haggard, 2010, but see Medina & Duckett, 2017 for alternative interpretation of the task), showing that the hand was perceived as even wider in congenital blindness. On the other hand, perceived hand shape in terms of how widened the hand was perceived (i.e. anisotropy) did not differ between congenital blindness and typical controls, whether measured by absolute tactile distance estimate (Giraud et al., 2025) or tactile distance comparison (Shahzad et al., 2024). Notably, these studies primarily focused on anisotropy in hand shape perception, whereby the hand is perceived as more widened than its veridical shape. This perceptual bias may be related to cortical neural organization such that the receptive fields of the hand and arm are elliptical with the long axis along the proximodistal axis of these body parts (Longo & Haggard, 2011). More broadly, this paradigm may be particularly sensitive to intrinsic properties of early somatosensory representations (Tamè et al., 2021), while providing limited dissociation between primary somatosensory processing and higher-order body form representations. Consequently, it may not fully capture the contribution of higher-order body representations to tactile distance perception. Therefore, whether and how early visual experience contributes to higher-order representations of body form remains an open question. The present study further investigates this question by examining tactile distance perception across multiple body regions in congenitally blind, late-blind, and sighted individuals.

Here we addressed this question in blindness using the same paradigm as in Taylor-Clarke et al. (2004), where individuals reported which of the two tactile distances applied to different skin areas felt longer. This paradigm has two main advantages. First, its error pattern reflects a dissociation between the intrinsic somatosensory system in S1, demonstrating a correcting effect from S1 cortical layout to the less distorted perception by the body form representation. Second, the task involves purely tactile distance perception without referring to additional spatial processes that may differ between sighted and blindness. The critical comparison involves the forearm and finger. If lifelong visual information is required for constructing the body form representation, the same tactile distance would be perceived longer on the finger than the forearm, and importantly to a greater extent in congenital blindness than controls. Alternatively, if a near-veridical body form representation can be constructed from non-visual sensory and motor experience, e.g. from excessive hand-object interaction, no difference would be observed between congenital blindness and controls.

## Methods

### Participants

Twenty-four individuals with blindness participated in the study. Participants with blindness were divided into two groups according to the onset of blindness and their reported visual memories. The early-blind (EB) group comprised individuals who became blind by the age of 2 and reported no visual memories, whereas the late-blind (LB) group comprised individuals who became blind later and reported retaining visual memories. Eleven participants with early blindness were recruited (ages: 30.5 ± 7.03, 1 female), with blindness onset ranging from birth to three years. See **Table 1** for complete information. Thirteen participants with late blindness were recuited (ages: 34.8 ± 8.9, 6 females), with blindness onset ranging from seven to 29 years. Twenty-six typical sighted controls (SC) were recruited (ages: 27.5 ± 6.76, 18 females). One SC participant was excluded because their accuracy on catch trials was below chance level (45.8%; see Experimental Design and Procedures). All studies were approved by the Institutional Review Board of the Institute of Neuroscience, Chinese Academy of Sciences and of Shanghai Jiao Tong University. Written informed consent was obtained from sighted participants and from the parents or legal guardians of minor participants. For participants with blindness, the informed consent form was read aloud by the experimenter. Their consent was then documented either by their fingerprint or, when applicable, by the signature of an accompanying person.

**Table 1.** Demographic and Clinical Characteristics of Blind Participants.

| Participant | Age | Group | Age at onset of visual impairment | Reported cause of visual impairment |
|---|---|---|---|---|
| B01 | 23 | LB | 12 years | Not reported |
| B02 | 36 | LB | 13 years | Not reported |
| B03 | 41 | EB | 1–2 years | High fever |
| B04 | 28 | EB | 3 months | Keratitis |
| B05 | 27 | EB | Approximately 1.5 years | Fall and subsequent diarrheal illness |
| B06 | 43 | EB | 3 years | Congenital retinitis pigmentosa and optic atrophy |
| B07 | 22 | EB | From birth | Progressive atrophy of the eyes during early childhood |
| B09 | 34 | LB | In the teens | Optic atrophy |
| B10 | 35 | LB | In the teens | Optic atrophy and glaucoma |
| B11 | 25 | EB | 1–2 years | Medical accident |
| B12 | 26 | EB | Very early childhood | Not reported |
| B13 | 56 | LB | 29 years | Blast-related ocular injury |
| B14 | 23 | LB | 7 years | Retinal detachment |
| B15 | 41 | LB | 15 years | Retinitis pigmentosa |
| B16 | 34 | EB | Childhood | Unspecified illness |
| B17 | 40 | LB | In the twenties | Retinal disease |
| B18 | 33 | EB | From birth | Unspecified congenital condition |
| B19 | 38 | LB | Primary school age | Retinitis pigmentosa |
| B20 | 24 | LB | 18 years | Optic atrophy |
| B21 | 32 | LB | 26 years | Brain tumor-induced optic nerve compression |
| B22 | 33 | LB | 18 years | Optic atrophy |
| B23 | 34 | EB | From birth | Unspecified congenital condition |
| B24 | 23 | EB | From birth | Congenital cataracts |
| B25 | 38 | LB | Childhood (night blindness) | Genetic mutation affecting the retina |

### Experimental design and procedures

In each trial, the participant was stimulated on two different skin areas with two points of a fixed distance and reported on which area they felt the distance between the two points longer. Skin areas included right index finger, right forearm (glabrous side), face, and forehead, for which all pairwise conditions were included. As with Taylor-Clarke et al., (2004), the core comparison involved index finger and forearm, with face a control area that matches the receptor density and perceived tactile distance with index finger. Forehead, with similar receptor density with face (Won et al., 2017), was included to examine whether the observed perceptual patterns were consistent across different facial regions. Additionally, left index finger was included and was paired only with the right index finger as a homologous control. Across trials, the distance between the two points was either 45mm or 55mm. Each of the 14 unique conditions (seven skin area pairs and two distances) was repeated 16 times, with the order of the stimulated skin area counterbalanced. Twenty-four catch trials in which skin areas were stimulated with different distances (45 mm and 55 mm respectively) were interspersed, resulting in 248 trials in total broken down into four blocks.

During the experiment, the participant placed their forearms and hands on the table, palm facing up and fingers opened flat. The two points were aligned with the proximodistal axis of the index finger and forearm, horizontally across the forehead, and along the angle of the jaw on the face. Typical controls were tested blindfolded. One blind participant could perceive blurry large-scale contours and was also blindfolded. The experiment was programmed in Matlab (R2025a; The MathWorks, Inc.). Trial orders were randomized for each individual and the experimenter input the participant’s verbal response from each trial to the program.

### Data analysis

As with Taylor-Clarke et al., (2004), our core comparison focuses on the right index finger and the forearm. For each participant, we calculated the proportion of trials in which the right index finger was judged to have a longer tactile distance than the right forearm. One-sample t-tests against 0.5 were conducted separately for each group to assess whether judgments showed a systematic bias toward either skin area. Control analyses were conducted between skin areas with similar receptor density and were expected to match in perceived tactile distance, i.e. between the right index finger and left index finger, between the right index finger and face, and between face and forehead.

The critical analysis concerned the difference between SC and EB. For each pair of skin areas, a Welch’s two-sample *t*-test was performed between SC and EB, implemented in R (version 4.6.1). Given the broad range of blindness onset ages in the LB group, which spanned childhood and adulthood, we additionally conducted exploratory analyses to examine potential individual differences associated with the timing of visual loss. These analyses were intended to complement the primary comparison between the SC and EB groups.

The analyses above only provides local comparison between each pair of skin area. To derive a global relative scale across skin areas, we fitted each participants’ trialwise responses with a Bradley-Terry model (Bradley & Terry, 1952; Usami 2009; Tallon et al., 2022; Wu et al., 2022). The model assumes that each item (i.e. skin area) has its hidden value (i.e. represented size) that cannot be directly observed, and that each pairwise comparison provides an observation with noise about the relative values of the two items. Specifically, the probability of choosing one skin area over another is modeled as a logistic function of the difference between their latent values:

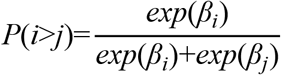

where β_i_ and β_j_ represent the latent values of skin areas *i* and *j*, respectively. The difference between two coefficients therefore represents the log odds ratio of choosing one area over the other:

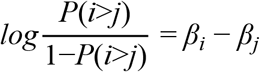

These coefficients provide a common latent scale across skin areas by jointly modeling all pairwise comparisons. For each participant, the hidden value of the right arm was fixed to zero as a reference, and the values of the other skin areas were estimated relative to this reference. The left index finger was not included because it was only paired with the right index finger. The resulting coefficients were then fitted into a linear-mixed model with Group (SC, EB) and Area (right index finger, face, forehead) as fixed factors and participants as a random factor, using the lmer function in R (version 4.6.1). An additional model including LBs was performed (3 Groups by 3 Areas) to examine the overall patterns across all groups.

## Results

### Pairwise comparisons of tactile distance across skin areas

Consistent with previous findings, sighted controls (SCs) perceived a given distance as longer on the right index finger than on the forearm (**Figure 1A**; P(Index finger) = 0.713, SD = 0.181, t(24) = 5.86, p < .001, 95% CI [0.638, 0.787], Cohen’s *d* = 1.17). No significant preference was observed in comparisons between the right index finger and face (P(Index finger) = 0.548, SD = 0.180, t(24) = 1.32, p = .200, 95% CI [0.473, 0.622], Cohen’s *d* = 0.264), or between the forehead and face (P(forehead) = 0.493, SD = 0.193, t(24) = -0.194, p = .848, 95% CI [0.413, 0.572], Cohen’s *d* = -0.03). These findings are consistent with the absence of a systematic perceptual difference between these pairs of skin areas reported in previous work (Taylor-Clarke et al., 2004). Interestingly, a given distance was perceived as longer on the left index finger more frequently than on the right index finger (P(Left Index finger) = 0.676, SD = 0.151, t(24) = 5.84, p < .001, 95% CI [0.614, 0.739], Cohen’s *d* = 1.17). One possibility is that the right index was stimulated overall more than the left index finger, potentially leading to habituation and a reduction in perceived tactile distance.

**Figure 1.**
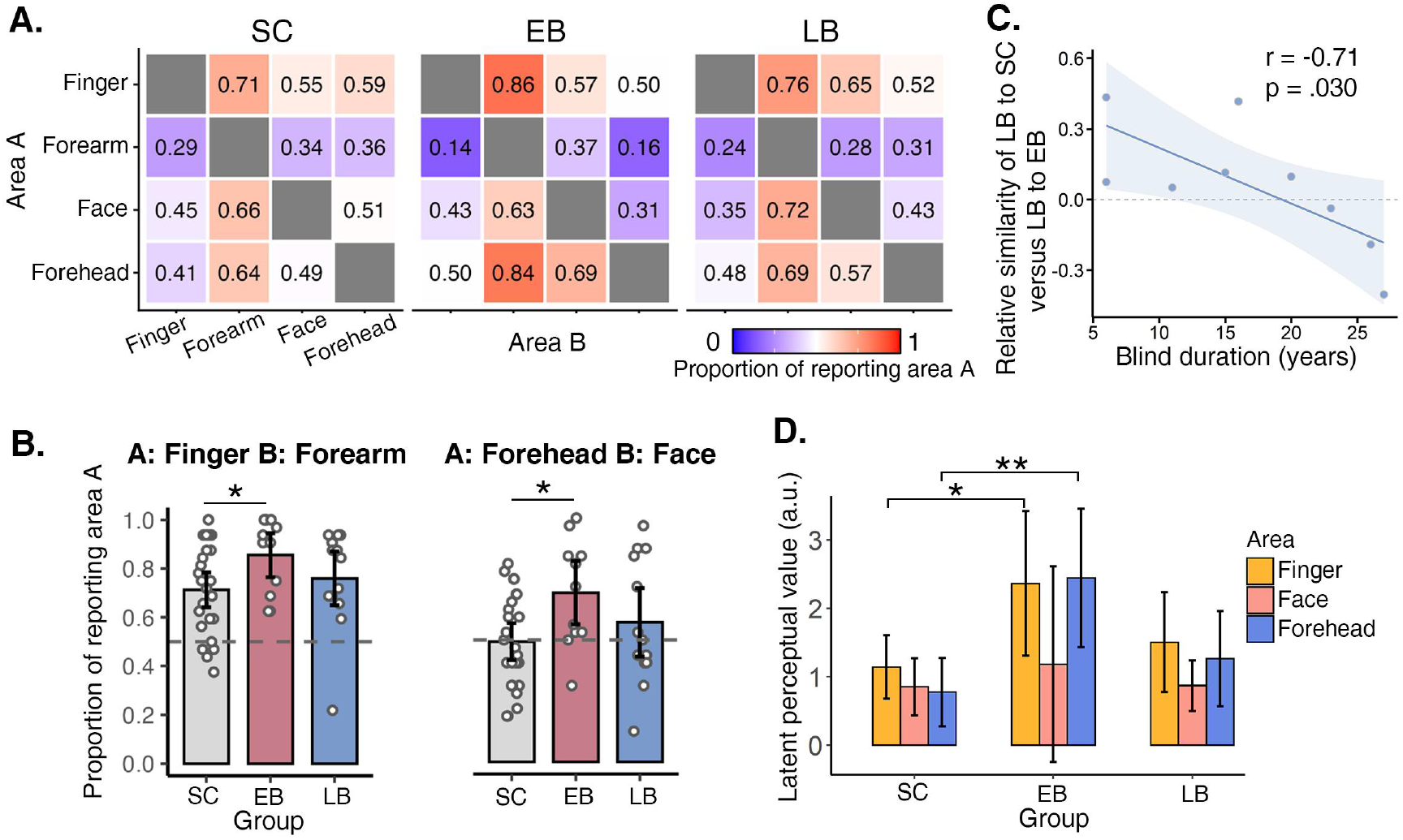
Summary of the main results. **(A)** Average proportion of trials in which area A was reported as having a longer tactile distance across sighted controls (SC), early blindness (EB), and late blindness (LB). Qualitatively, tactile distances on the finger (i.e. right index finger) and forehead were more likely to be perceived as longer than those on other body areas, particularly in EB. Unless otherwise noted, finger refers to the right index finger. **(B)** Porportion of trials in which the right index finger was reported when paired with the forearm, and the forehead was reported when paired with the face. Tactile distances were more likely to be perceived as longer on the index finger than on the forearm, and on the forehead than on the face, in EB, with both effects being more pronounced than in SC. **(C)** In LB, longer blindness duration was associated with a relative shift in behavioral similarity away from SC and toward EB participants **(D)** Results from the Bradley–Terry model. The latent size of each body area was estimated relative to the forearm, which served as the reference. Consistent with the raw response proportions, the index finger and forehead were perceived as longer in EB than in SC.

Our core comparison focuses on the difference between SCs and EBs regarding the relative perceived distance between the right index finger and forearm. EBs also perceived a given distance as longer on the index finger than on the forearm (P(Index finger) = 0.855, SD = 0.153, t(10) = 7.72, p < .001, 95% CI [0.753,0.958]; **Figure 1A** and **1B**). Importantly, this preference was significantly stronger in EBs than in SCs (Group difference = 0.142, t(22.6) = 2.43, p = .023, 95% CI [0.021, 0.264], Cohen’s d = 0.823; **Figure 1A and 1B**). These findings suggest that the distortion between the index finger and forearm was magnified in individuals who were blind since early life relative to SCs. Late-blind participants (LBs) also showed a preference for judging the index finger as longer than the forearm (P (Index finger) = 0.760, SD = 0.203, t(12) = 4.62, p < .001, 95% CI [0.637, 0.882]). However, their performance did not differ significantly from either SCs (t(22.1) = 0.704, p = .489, 95% CI [-0.092, 0.186], Cohen’s d = 0.250) or EBs (t(21.8) = 1.31, p = .202, 95% CI [-0.055, 0.246], Cohen’s d = 0.526).

An additional finding concerned the comparison between the forehead and face. Whereas SCs showed no systematic preference between these two areas, EBs were more likely to perceive a given distance as longer on the forehead than on the face (*P*(forehead) = 0.693, SD = 0.220, *t*(10) = 2.92, *p* = .015, 95% CI [0.546, 0.841], Cohen’s *d* = 0.880). This preference was also significantly stronger in EBs than in SCs (mean difference = 0.201, *t*(17.15) = 2.62, *p* = .018, Cohen’s *d* = 0.996; **Figure 1A and 1B**). LBs, in contrast, showed no significant preference for either area (P(forehead) = 0.572, SD = 0.258, *t*(12) = 1.01, *p* = .334, 95% CI [0.416, 0.728], Cohen’s *d* = 0.279). This result suggests that the relative perception of tactile distance on the forehead and face differed between EBs and SCs.

### Behavioral similarity within the late-blind group

Because age of blindness onset in the LB group ranged from childhood (7 years) to adulthood (29 years), this group may encompass substantial variability in visual experience during development. We therefore explored whether individual differences in LB participants’ behavioral profiles were associated with age of blindness onset or duration of blindness. Specifically, we considered the possibility that participants with earlier blindness onset might show behavioral profiles more similar to those of EBs, whereas those with later onset might show profiles more similar to SCs.

For each participant, we defined a behavioral profile as the set of proportions of trials on which a given skin area was judged to have the longer tactile distance in each unique pairwise comparison. These values corresponded to the lower triangular portion of the matrices shown in **Figure 1A**; comparisons involving the left index finger were excluded. For each LB participant, we calculated a relative behavioral similarity score by subtracting the correlation between their behavioral profile and the EB group-average profile from the correlation between their profile and the SC group-average profile. We then examined the association between this score and both age of blindness onset and blindness duration.

Age of blindness onset was not significantly associated with relative behavioral similarity (Pearson’s correlation coefficient *r* = −0.327, *p* = .390, 95% CI [−0.814, 0.430]). In contrast, blindness duration showed a significant negative correlation with the similarity score (*r* = −0.710, *p* = .030, 95% CI [−0.935, −0.099]; **Figure 1C**). This association remained significant after statistically controlling for age of blindness onset (partial *r* = −0.749, *p* = .032). Thus, individual differences in blindness duration, rather than age of onset alone, were associated with variation in the behavioral profiles of LB participants.

### Bradley-Terry estimates of relative perceptual scales

The pairwise comparisons described above provide local measures of perceived tactile distance between specific pairs of skin areas. To estimate a global relative perceptual scale across skin areas, we additionally fitted a Bradley–Terry model to each participant’s trial-wise binary choices (**Figure 1D**). The model yielded a latent perceptual value for each skin area, with the forearm serving as the reference area (β = 0). Higher values indicate a greater likelihood that a given area was judged to have a longer tactile distance than the forearm.

We first compared SCs and EBs using a linear mixed-effects model with Group (SC, EB) and Area (right index finger, forehead, face) as fixed factors and participant as a random factor. The analysis revealed a main effect of Group (F(1, 34) = 5.76, p = .022, partial ƞ^2^ = 0.14), driven by an overall larger latent value in EBs than SCs. The main effect of Area was also significant (F(2, 68) = 4.95, p = .010, partial ƞ^2^ = 0.13) as driven by a larger latent value of the index finger than face (Tukey adjusted p = .011, Cohen’s d = 0.760) and a marginally larger latent value of forehead than face (Tukey adjusted p = .051, Cohen’s d = 0.611). Importantly, there was a significant interaction between Group and Area (F(2, 68) = 3.76, p = .028, partial ƞ^2^ = 0.10), such that EBs showed a larger latent value than controls in the index finger (Tukey adjusted p = .025, Cohen’s d = 1.26) and forehead (Tukey adjusted p = .003, Cohen’s d = 1.72), but not in face (Tukey adjusted p = .534, Cohen’s d = 0.343). These results indicate that, relative to the forearm, the right index finger and forehead had higher latent perceptual values in EBs than in SCs.

We next included LBs in the analysis to examine the overall pattern across all three groups. A linear mixed-effects model with Group (SC, EB, LB) and Area (right index finger, forehead, face) as fixed factors and participant as a random factor revealed significant main effects of Group (*F*(2, 46) = 3.27, *p* = .047, partial η² = 0.12) and Area (*F*(2, 92) = 6.60, *p* = .002, partial η² = 0.13), as well as a nonsignificant Group × Area interaction (*F*(4, 92) = 2.02, *p* = .098, partial η² = 0.08). Post-hoc comparisons for Group indicated that the overall group effect was driven by a difference between SCs and EBs (Tukey-adjusted *p* = .037, Cohen’s *d* = 1.15), whereas neither the SC–LB comparison (Tukey-adjusted *p* = .748, Cohen’s *d* = 0.309) nor the EB–LB comparison (Tukey-adjusted *p* = .236, Cohen’s *d* = 0.838) was significant. The main effect of Area reflected lower latent values for the face than for both the right index finger (Tukey-adjusted *p* = .002, Cohen’s *d* = 0.751) and the forehead (Tukey-adjusted *p* = .028, Cohen’s *d* = 0.562). No significant difference was found between the right index finger and forehead (Tukey-adjusted *p* = .657, Cohen’s *d* = 0.188).

## Discussion

The present study investigated whether early visual experience contributes to the development of a relatively veridical representation of body form, as reflected in perceived tactile distances across different skin areas. We found that individuals with early blindness (EB) were more likely than typically sighted controls (SCs) to perceive a given tactile distance as longer on the index finger than on the forearm. This difference was evident in both the direct pairwise comparisons and the global relative perceptual scales derived from the Bradley–Terry model. In SCs, no systematic differences were observed in comparisons between the index finger and face or between the forehead and face. Together, these findings suggest that early visual experience contributes to the calibration of perceived tactile distance across body regions, although the precise representational mechanisms underlying this effect remain to be established.

Previous studies have investigated the role of visual experience in body representation, but their findings have not been entirely consistent. Kottu and Lazar (2025), using a hand landmark localization task, reported greater distortions in the implicit representation of hand shape in individuals with early blindness than in sighted controls. However, the interpretation of landmark localization performance as a direct measure of body representation has been questioned. Medina and Duckett (2017), for example, demonstrated that apparent distortions in hand landmark localization can arise, at least in part, from domain-general biases in spatial localization. The present study provides complementary evidence using a different behavioral approach. Rather than requiring participants to reconstruct the locations of anatomical landmarks, our task assessed relative tactile distance directly across skin areas. Although this task still involves spatial judgments, it does not require explicit localization of body landmarks or reconstruction of hand shape. The observed group differences therefore extend previous findings by showing that early visual experience is also associated with the relative scaling of tactile distances across body regions.

Our findings also differ from recent studies examining tactile distance anisotropy within individual body parts. Giraud et al. (2025) reported broadly similar patterns of tactile hand-space distortion in blind and sighted participants, suggesting that typical distortions of tactile space can persist in the absence of visual experience. Similarly, Shahzad et al. (2025) found tactile distance anisotropies in both early-blind and sighted individuals, although the magnitude of the anisotropy was reduced in the forearm of blind participants. These findings do not necessarily contradict the present results, as the paradigms probe different aspects of tactile spatial representation. Tactile anisotropy concerns the perceived relative distances along different axes within a single body part and has neural correlates in S1 (Tamè et al., 2021), whereas our primary comparison concerns the relative scaling of tactile distances across different body parts that partially reflects rescaling beyond S1 representations. It is possible that these forms of distortion depend, at least in part, on different representational processes and are differentially influenced by visual experience. Thus, visual experience may contribute to calibrating the relative tactile scale across body regions without being necessary for the emergence of all distortions in the representation of individual body parts.

An alternative explanation for our findings is that differences in primary somatosensory processing, rather than higher-order body representation, contribute to the enhanced index-finger–forearm difference in EB. Tactile distance perception is influenced by the organization of early somatosensory representations, and experience-dependent plasticity in primary somatosensory cortex (S1) has been demonstrated following extensive sensorimotor training or repeated tactile stimulation (Elbert et al., 1995; Jenkins et al., 1990; Recanzone et al., 1992). Given the increased reliance on manual exploration in blindness, it is therefore possible that differences in tactile experience contribute to changes in S1 processing. However, evidence that increased tactile use necessarily leads to a more extensive or reorganized S1 hand representation in blindness is mixed. For example, a 7-T fMRI study of individuals with congenital anophthalmia found no clear evidence of more pronounced finger representations in S1, despite their superior tactile acuity (Wesselink et al., 2021). Moreover, enhanced tactile acuity in blindness appears to be particularly associated with the fingers used for Braille reading, rather than being uniformly distributed across the fingers (Van Boven et al., 2000; Wong et al., 2011). None of the blind participants in the present study were Braille readers, making this particular form of use-dependent S1 plasticity less likely to account for our findings.

In contrast to early blindness, performance of late blindness showed patterns that overall more resembled sighted controls. This may suggest that higher-order body form representation undergoes an early sensitive period during which visual experience contributes to their establishment and calibration, and after which they become relatively stable and less dependent on continuous visual input. On the hand, the late blind group in our study included individuals with blind onset during childhood, as early as seven years old. A coherent body form representation relies on multisensory integration (Pavani & Zampini, 2007; Giurgola et al., 2022), a process that continues to develop throughout childhood (Cowie et al., 2016; Roder et al., 2013). Moreover, representation of body size continuously update during pubertal growth (de Haan et al., 2018). Therefore, visual loss during childhood or adolescence may interfere with the maturation and updating of higher-order body representations, whereas visual loss in adulthood may have less impact once these representations are relatively established. Importantly, our exploratory analysis of individual differences within the LB group revealed that behavioral similarity to the sighted and early-blind groups was associated with blindness duration, but not significantly with age of blindness onset. Specifically, longer blindness duration was associated with greater behavioral similarity to the early-blind group, and this association remained significant after controlling for age of blindness onset. This finding raises the possibility that even after visual experience has contributed to the initial establishment of body representations, these representations may remain susceptible to changes associated with prolonged visual deprivation. The overall intermediate pattern observed in the LB group in the Bradley–Terry analysis may therefore reflect heterogeneous influences of both developmental timing and subsequent experience without necessarily indicating a uniform intermediate stage between early blindness and sightedness. Nevertheless, given the relatively small sample size and the exploratory nature of this analysis, these interpretations should be treated with caution. Future studies with larger samples and a broader range of blindness onset ages and durations are needed to disentangle the contributions of developmental timing and post-onset experience to the organization of higher-order body form representations.

In addition to the index finger, tactile distance on the forehead was perceived as longer in early blindness and to a greater extent than in sighted controls. This effect was specific to the forehead and was not observed on the face. To our knowledge, few previous studies have specifically examined forehead tactile spatial perception in blindness, and the basis of this finding remains unclear. Unlike the fingers, the forehead is not typically subject to extensive tactile exploration or specialized use in daily activities, especially not relative to face, hence unlikely showing use-dependent plasticity. One potentially relevant anatomical distinction between the forehead and the face is their peripheral sensory innervation: forehead sensation is primarily conveyed through the ophthalmic division (V1) of the trigeminal nerve, which also delievers somatosensory information from the ocular region, whereas face sensation is conveyed primarily through the maxillary division (V2; Leston, 2009; Kim et al., 2024). However, this anatomical overlap does not provide a direct explanation for the present finding, as blindness involves the loss of visual input rather than the loss of ocular somatosensory input. Whether the observed forehead-specific difference reflects changes in peripheral or central sensory processing, or higher-order body representations, remains to be determined. Future research with more detailed behavioral assessments combined with neuroimaging measures will be needed to examine the underlying mechanisms.

Our study has several limitations. First, the relatively modest sample size, particularly in the blind groups, limited statistical power for more fine-grained analyses, including examining individual differences in performance and their relationships with the timing and duration of blindness. The cross-sectional design also precludes establishing the developmental trajectory of body form representations. Second, the present study examined a limited set of body regions, and whether the observed patterns generalize to other body parts, particularly those differing in their accessibility to manual exploration and involvement in everyday movements, remains to be investigated. Future studies with larger samples and longitudinal designs are needed to better characterize individual differences in body form representations, establish their developmental trajectories, and examine how the timing and duration of visual deprivation relate to perceptual outcomes.

## Acknowledgements

We sincerely thank all participants with blindness and their families for their participation in this study. This work was supported by the National Science and Technology Innovation 2030 Major Project 2021ZD0204200 (2021ZD0204204) to Y. L.

